# Antimicrobial Susceptibility Testing of *Chlamydophila pneumoniae*: Pilot Study for Alternative Methods that Address the Complex Chlamydial Life Cycle

**DOI:** 10.1101/403295

**Authors:** Laura S. Stewart, Sandra Alvarez-Macias, Rendie McHenry, James D. Chappell, Charles W. Stratton

## Abstract

Conventional susceptibility testing of *Chlamydophila pneumoniae* does not account for the complex life cycle that takes place in an obligatory intracellular niche and involves multiple morphological forms. Some of these forms (elementary bodies and cryptic bodies) cause persistence and are not susceptible to antimicrobial agents. Therefore, we describe a pilot study for alternative methods of susceptibility testing of *C. pneumoniae*. These methods include delaying the addition of antimicrobial agents to cell cultures for 48 hours to allow the development of an established chlamydial infection as well as the use of reverse transcriptase quantitative PCR [RT-qPCR] to measure messenger RNA. Using these methods, susceptibility testing of an established infection in Hep2 cells was compared with conventional susceptibility testing of *C. pneumoniae*.

Conventional antimicrobial susceptibility testing of *C. pneumoniae* results in MICs and MBCs that would suggest that recommended treatment regimens of 2 to 3 weeks of antibiotics such as doxycycline, clarithromycin, levofloxacin, and rifabutin would be sufficient. However, susceptibility testing using an established chlamydial infection in HEp2 cells reveals that none of these agents are as inhibitory and/or bactericidal as indicated by conventional methods. The results of this pilot study suggest that further evaluation of chlamydial susceptibility testing methods using established infection in cell cultures and RT-qPCR to measure messenger RNA are needed to optimize treatment recommendations for chronic *C. pneumoniae* infections.

## Introduction

Conventional susceptibility testing of *C. pneumoniae* is done by inoculating cell cultures with infectious elementary bodies, centrifuging for 30-60 minutes, and then adding the antimicrobial agent to be tested (1-4). Following 3 days of incubation, cell cultures are stained with a genus-specific fluorescent monoclonal antibody stain and examined for chlamydial inclusions. The MIC is defined as the lowest concentration of drug that inhibits the development of inclusions (1-4).

This approach evaluates the initial infection of cell cultures by *C. pneumoniae* infectious elementary bodies, but may not sufficiently assess the unique chlamydial life cycle. Once the intracellular chlamydial infection has been established for several days, the infected cells will contain non-replicating cryptic bodies and condensing elementary bodies as well as replicating reticulate bodies (5). These cryptic bodies and elementary bodies generally are not susceptible to antimicrobial agents, yet are responsible for the persistence of *C. pneumoniae* (5-8). This chlamydial persistence often results in the inability of antimicrobial therapy to eradicate established *C. pneumoniae* infection and is thought to be clinically important (6-8). For example, persistent respiratory tract infections by *C. pneumoniae* have been described; repeated antimicrobial therapy does not seem to be able to eradicate this pathogen in these patients (9-11).

Because conventional susceptibility testing of *C. pneumoniae* may not account for the complex chlamydial life cycle leading to persistence, we describe a pilot study that investigates alternative methods for susceptibility testing of *C. pneumoniae*. These alternative methods include delaying the addition of antimicrobial agents to the cell culture for 48 hours to allow the development of an established infection, the use of reverse transcriptase quantitative PCR [RT-qPCR] to measure chlamydial messenger RNA for major outer-membrane protein [MOMP] and heat-shock protein 60 [HSP60], and an infectivity assay to detect the presence of elementary bodies after antimicrobial therapy. RT-qPCR has been described as a useful method for susceptibility testing of *Chlamydiae* by numerous investigators (12-15). These alternative methods of susceptibility testing were compared with conventional susceptibility testing of *C. pneumoniae*, which assesses only the initial infection of HEp2 cells by elementary bodies.

## Results

### Evaluation of initial infection of HEp2 cells with *C. pneumoniae* elementary bodies

Initial infection of HEp2 cells with *C. pneumoniae* elementary bodies was evaluated for 5 days following infection using multiple methods for detection of persistence. Expression of HSP60 and MOMP genes in infected cell culture lysates was measured daily using RT-qPCR [Fig 1A]. These targets were chosen because HSP60 and MOMP expression is often differentially regulated in acute versus chronic infection (6). Both HSP60 and MOMP expression rose logarithmically for the first 2 days following infection. MOMP plateaued at around a 100 fold increase over input while HSP60 expression underwent a further modest increase between days 3 and 5. Infectivity of *C. pneumoniae* in cell cultures was measured by passing infected cell lysates onto fresh HEp2 cells [Fig 1B]. Following 1 day of infection, infectivity was just barely detected at 2.75 log_10_ inclusion-forming units [IFU] per ml, corresponding to a 1.55 log declined from the day 0 level [input].

**Fig 1.**
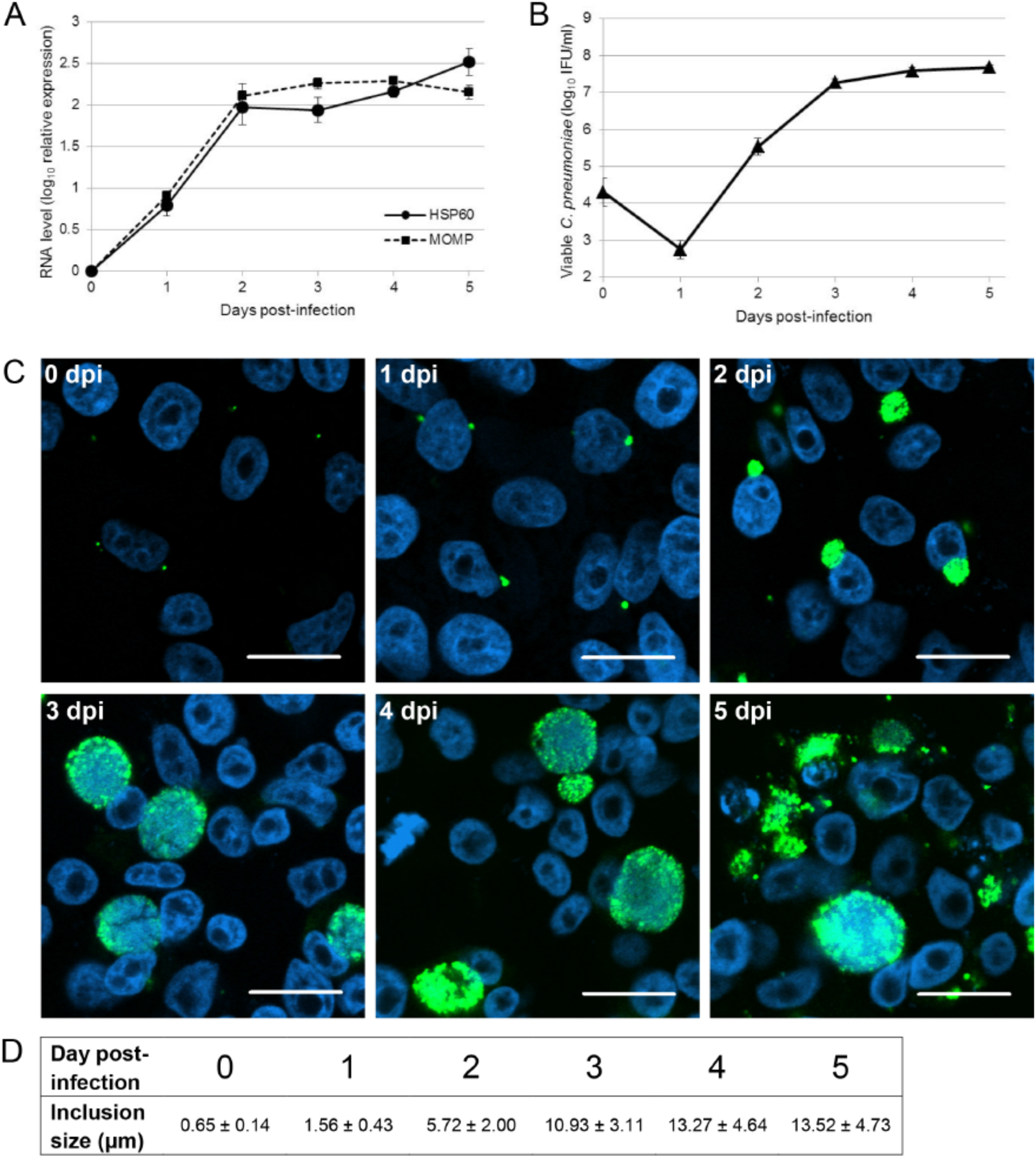
Time course after initial infection of HEp2 cells with *C. pneumoniae* elementary bodies. HEp2 cells were infected at an MOI of 0.2 IFU/cell and incubated for 3 days. [A] Expression of HSP60 and MOMP mRNA was measured ininfected cell lysates using RT-qPCR. Results were normalized to RNAseP expression and are plotted relative to samples collected at day 0 post-infection.[B] Infectivity titer was measured in cell lysates. [C] Infected cells were fixed, stained with an anti-cHSP60 antibody [green] and ToPro3 [blue, DNA], and imaged using confocal microscopy. Scale bars, 20 μm. [D] Average inclusion size [+/- SD] was measured from confocal images of infected cells.

Infectivity increased approximately 100 fold over the subsequent two days and then plateaued at day 4 post-infection. Inclusion morphology was monitored over the time course of infection using confocal immunofluorescence microscopy [Fig 1C], and diameters of inclusions were measured on representative images [Fig 1D]. Cells were infected, fixed at various intervals, and stained with anti-HSP60antisera. At day 0 post-infection [fixed immediately following inoculation], small green dots of fluorescence were observed. This staining presumably represents attached and/or internalized EBs. At day 1 post-infection, small, brightly stained inclusions were present. Between day 1 and 2 post-infection, inclusion diameter [mean ± SD] increased from 1.56 ± 0.43 μm to 5.72 ± 2.00 μm. By day 3 post-infection, large inclusions [10.93 ± 3.11 μm] were observed. The staining of these inclusions was generally more punctate in comparison to day 2 post-infection. As the infection progressed, the range of inclusion size increased and is reflected in the increasing standard deviation of diameter measurement over time. Inclusion size increased further at days 4 and 5 post-infection [13.27 ± 4.64 μm and 13.52 ± 4.73 μm, respectively]; however, while large inclusions were still observed at day 5, many inclusions had lysed by that time, leaving a significant amount of debris.

### Assessment of antimicrobial treatment of HEp2 cells after initial infection by *C. pneumoniae* elementary bodies using conventional methodology

The susceptibility of *C. pneumoniae* to many antimicrobial agents has been evaluated to determine the minimum inhibitory concentration [MIC] (1-4) and minimum bactericidal concentration [MBC] (16-19) of these agents. Methodologies used have varied by study. Although standardization of antimicrobial susceptibility testing methods for *C. pneumoniae* has been recommended (20), suchstandardization has not been accomplished. Most commonly, *C. pneumoniae* is added to cells, followed by a 30-60 minutes of centrifugation and then addition of serially diluted antibiotics. After 3 days of culture, inhibition of chlamydial replication is ascertained either by PCR or detection of inclusions by immunofluorescence. The lowest concentration of drug that inhibits replication is designated as the MIC. To determine the MBC, drug-treated culture lysates are passed onto fresh monolayers, and chlamydial replication is detected by PCR or immunofluorescence after 3 days of culture. For our pilot study, we chose to analyze representative drugs of the major classes of antimicrobial agents that penetrate eukaryotic cells and are known to inhibit *C. pneumoniae* replication: doxycycline [tetracycline], clarithromycin [macrolide], rifabutin [rifamycin], and levofloxacin [fluoroquinolone]. We used a well-characterized strain of *C. pneumoniae* [AR-39] that was initially isolated from a coronary atheroma (21). In Fig 2A, we determined MICs of these agents using RT-qPCR. HEp2 cells were infected at a multiplicity of infection [MOI] of 0.2. Following centrifugation, serial dilutions of doxycycline, clarithromycin, rifabutin, and levofloxacin were added to cultures. HSP60 and MOMP mRNA expression normalized to RNAseP expression was measured following 3 days of incubation and monitored relative to day 0 post-infection. A day 3/day 0 ratio less than 1, or less than input, was considered evidence of inhibited replication. Full inhibition of replication was observed following treatment with 0.25 μg/ml doxycycline, 0.016 μg/ml clarithromycin,0.016 μg/ml rifabutin, and 5 μg/ml levofloxacin. These values are comparable to published MICs for these drugs (1-4,16-19). HSP60 and MOMP mRNA levels were similar in uninhibited infection; however, MOMP levels were consistently less than HSP60 when inhibitory concentrations of all antimicrobial agents and sub-inhibitory concentrations of doxycycline, clarithromycin, and levofloxacin were used. In levofloxacin-treated cells, this difference affected the determination of the MIC as 1 μg/ml and 5 μg/ml were needed to reduce MOMP and HSP60 expression, respectively, below input. The following concentrations were used in all subsequent experiments: 1 μg/ml doxycycline, 0.25 μg/ml clarithromycin, 0.25μg/ml rifabutin, and 1 μg/ml levofloxacin. With exception of levofloxacin, these concentrations are fully inhibitory (1-4,16-19) and are also achievable serum levels in humans (22-25). For levofloxacin, 1 μg/ml achieved 93% inhibition of HSP60 expression and full inhibition of MOMP expression and is a concentration that can be achieved using standard dosing recommendations (25).

**Fig 2.**
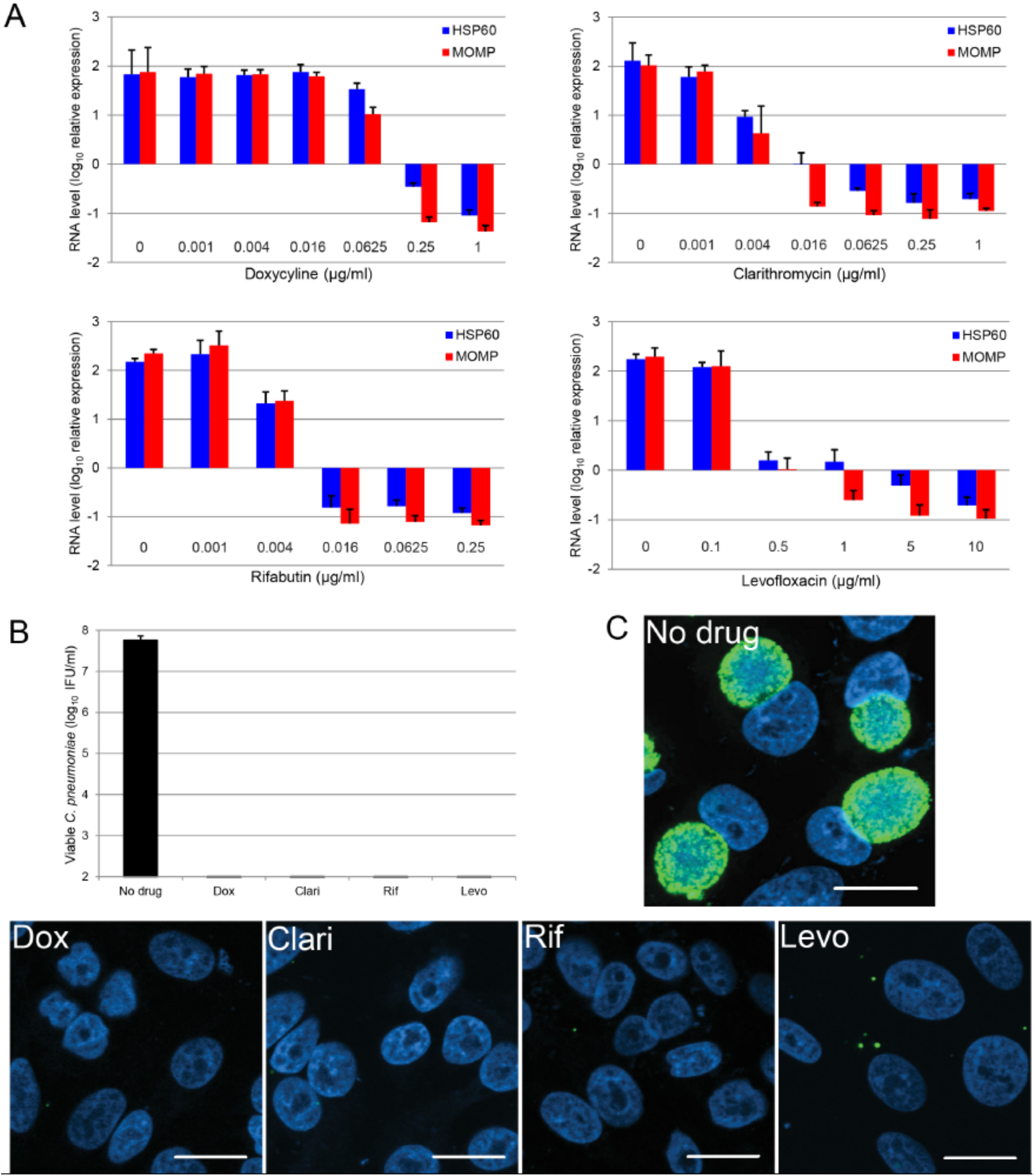
Time course for antimicrobial treatment of HEp2 cells after initial infection with *C. pneumoniae* elementary bodies. HEp2 cells were infected at an MOI of0.2 IFU/cell and treated with doxycycline, clarithromycin, rifabutin, or levofloxacinfor 3 days. [A] The minimum inhibitory concentration of each antibiotic was determined by treating cells with increasing concentrations of each agent. Expression of Hsp60 and MOMP mRNA was measured in infected cell lysates using RT-qPCR. Results were normalized to RNAseP expression and are plotted relative to samples collected at day 0 post-infection. [B, C] Infected cells were treated with 1 μg/ml doxycycline, 0.25 μg/ml clarithromycin, 0.25 μg/ml rifabutin, or 1 μg/ml levofloxacin. Following three days of treatment, infectivity titer was measured in cell lysates [B] or cells were stained with an anti-cHSP60 antibody [green] and ToPro3 [blue, DNA], and imaged using confocal microscopy [C]. Scale bars, 20 μm.

The infectivity of *C. pneumoniae* cultures following antimicrobial treatment was measured by passaging infected cell lysates onto fresh cells (Fig 2A). No titer was detected in any of the treated cultures. Confocal microscopy revealed that treatment prevented inclusion development (Fig 2C). In doxycycline-, clarithromycin-, and rifabutin-treated cultures, HSP60 staining was limited, and the few small fluorescent foci detected were similar to those observed in cultures fixed at day 0 post-infection. Inclusions in levofloxacin-treated cultures were slightly larger and more abundant than in the other antimicrobial-treated cultures (Fig 1D). However, with an average diameter of 0.81 ± 0.25 μm, these structures were smaller than those observed in an untreated infection at day one post-infection.

### Chlamydial infectivity following removal of antimicrobial agent

The results shown in Fig 2 provide evidence for bactericidal activity against *C. pneumoniae* for the antimicrobial agents tested and are consistent with published studies (3,17,19). However, *C. pneumoniae* may exist in a viable but non-infectious state (5-8). To ascertain whether viable *C. pneumoniae* remained after treatment with antimicrobial agents, cultures were initially incubated in the presence of doxycycline, clarithromycin, rifabutin, or levofloxacin for 1 or 3 days, followed by removal of antibiotic-containing medium, washing the cells, and then incubating cells in antimicrobial-free medium for 3 days. Both mRNA and infectivity were measured in cell culture lysates [Fig 3]. Removal of doxycycline led to significant recovery of mRNA expression and replication in a manner inversely related toduration of treatment. In clarithromycin-treated cell infections, a minimal amount of HSP60 mRNA above input was detected in samples treated for one day.

**Fig 3.**
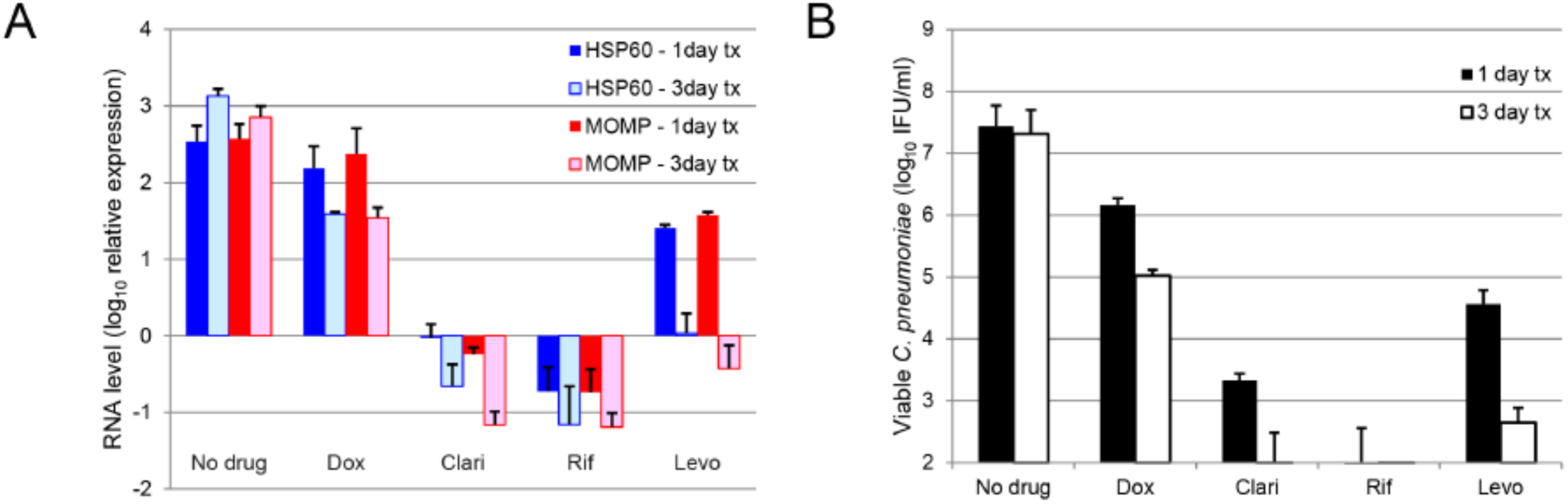
Evaluation of *C. pneumoniae* infectivity following antimicrobial treatment after initial infection of HEp2 cells by elementary bodies. HEp2 cells were infected at an MOI of 0.2 IFU/cell and treated with 1 μg/ml doxycycline, 0.25 μg/ml clarithromycin, 0.25 μg/ml rifabutin, or 1 μg/ml levofloxacin. After washingcells, culture medium was replaced with antimicrobial-free medium at either day 1 or day 3 post-infection and cultures were incubated for an additional three days. [A] Expression of HSP60 and MOMP mRNA was measured in infected cell lysates using RT-qPCR. Results were normalized to RNAseP expression and are plotted relative to samples collected at day 0 post-infection. [B] Infectivity titer was measured in infected cell lysates.

However, a low infectious titer was observed. Treatment for three days with clarithromycin reduced the mRNA level below input, and infectivity was virtually undetectable. The infection did not recover following either one or three days of rifabutin treatment as mRNA level was less than input, and infectious units were not detectable. Interestingly, 1 day of levofloxacin treatment did not prevent recovery of infection; however, 3 days of treatment nearly eliminated any recovery of mRNA expression or infectivity.

### Evaluation of antimicrobial treatment for cells with established infection by *C. pneumoniae*

In a natural infection, antimicrobial agents are not administrated immediately following exposure/infection with *C. pneumoniae*. To more closely mimic treatment of a clinical infection, we tested the *in vitro* response to antimicrobial treatment of an established infection by *C. pneumoniae*. HEp2 cells were initially infected with elementary bodies and then incubated in the absence of antimicrobial agents for 48 hours. At day 2 post-infection, doxycycline, clarithromycin, rifabutin, or levofloxacin was added to the infected cells. At day 4 post-infection, mRNA levels and *C. pneumoniae* infectivity were measured in culture lysates. Antimicrobial treatment was initiated at day 2 post-infectionbecause at this point in the infectious cycle, reticulate bodies are proliferating and some have transitioned to elementary bodies or to cryptic bodies (5-9).

Furthermore, the culture titer at day 2 was approximately 2.5 log_10_ IFU/ml higher than at day 1 post-infection, and further increased by approximately 2 log_10_ IFU/ml with two more days of incubation in the absence of antimicrobial agents [Fig 1B]. Treatment with either doxycycline or clarithromycin resulted in an approximately 1.5-2-fold decrease in HSP60 and MOMP levels at the same time point. Levofloxacin-treated infection resulted in a 2.7-fold reduction in HSP60 and 8.6-fold reduction in MOMP levels. The largest effect was seen in rifabutin-treated infection, with a 25- and 94-fold reduction in HSP60 and MOMP levels, respectively [Fig 4A]. A one-way ANOVA calculated on both HSP60 and MOMP mRNA levels confirmed statistical significance of declines in these targets [*P*<0.0001] [Fig 4B]. Of note, only clarithromycin-treated infection did not display significant reduction of either RNA levels or titer in comparison to day 2 treated infections; however, both parameters were significantly lower than an untreated infection at day 4 post-infection. Interestingly, despite differences in RNA levels, all treatment groups demonstrated equivalent infectious titers at day 4, which were unchanged relative to day 2 [Fig 4C, 4D]. Day 4 infectivity titers under eachantimicrobial-treatment condition were significantly lower than those achieved in untreated control culture [Fig 4D].

**Fig 4.**
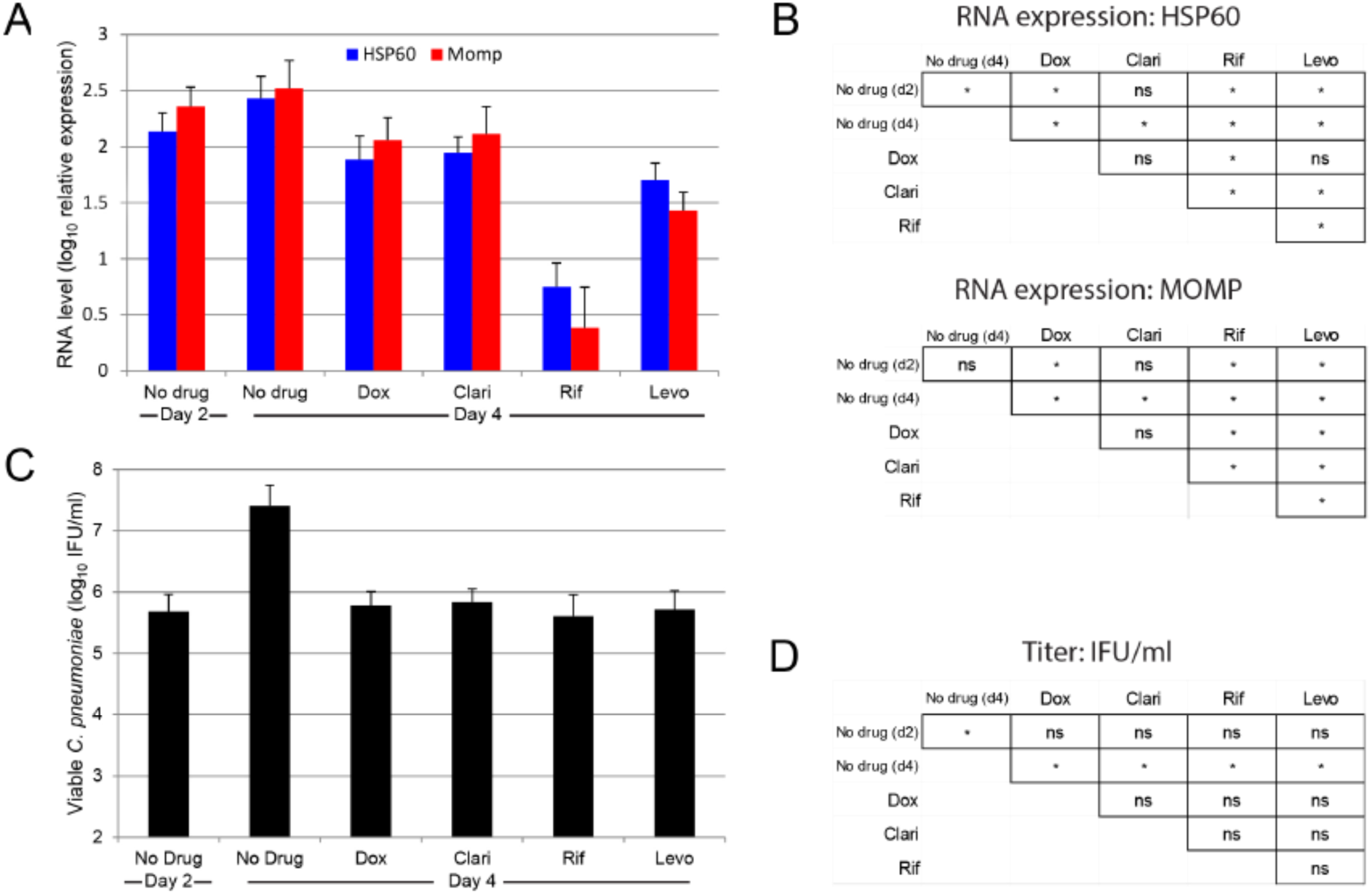
Evaluation of *C. pneumoniae* infectivity following antimicrobial treatment of established infection in HEp2 cells. HEp2 cells were infected at an MOI of 0.2 IFU/cell. Following two days of culture, 1 μg/ml doxycyline, 0.25 μg/ml clarithromycin, 0.25 μg/ml rifabutin, or 1 μg/ml levofloxacin was added and cellswere cultured for an additional two days. [A] Expression of HSP60 and MOMP mRNA was measured in infected cell lysates using RT-qPCR. Results were normalized to RNAseP expression and are plotted relative to samples collected at day 0 post-infection. [B] Infectivity titer was measured in infected cell lysates. [C,D] Results of Tukey’s multiple comparison test on mRNA level [C] and titer [D] data. ns, not significant; *, *P*<0.05.

### Replication recovery following antimicrobial treatment for cells with established infection with *C. pneumoniae*

To determine whether residual infectivity in antimicrobial-treated *C. pneumoniae* cultures is sufficient to reestablish normal bacterial replication, we monitored viable bacterial titers in antimicrobial-free medium for 6 days following treatment with antimicrobial agents. Infected HEp2-cell cultures were treated with antimicrobial agents from days 2-4 as described above, followed by cell-washing and antimicrobial-free medium exchange at days 4, 6, and 8. Samples were collected at days 2, 4, 6, 8, and 10 for quantification of *C. pneumoniae* mRNA and infectivity [Fig 5]. In doxycycline-treated infection, mRNA levels moderately decreased in the presence of the antimicrobial agent. Removal of doxycycline led to an increase in HSP60 and MOMP mRNA and an increase in titer at day 6 post-infection. The mRNA levels in clarithromycin-treated cultures remained steady throughout the infection, except for a drop in MOMP expression at day 6. Infectious titers remained stably low through day 8 post-infection and then increased markedly. Rifabutin treatment produced substantial declines in bacterial mRNA levels and infectivity, followed by rapid rebound after reaching a nadir. Infectivity recovery trailed restoration of mRNA expression byapproximately two days. In levofloxacin-treated infection, mRNA levels decreased through day 6 post-infection and then rose over the remaining course of infection. Similarly, infectivity increased between day 6 and 8 post-infection.

**Fig 5.**
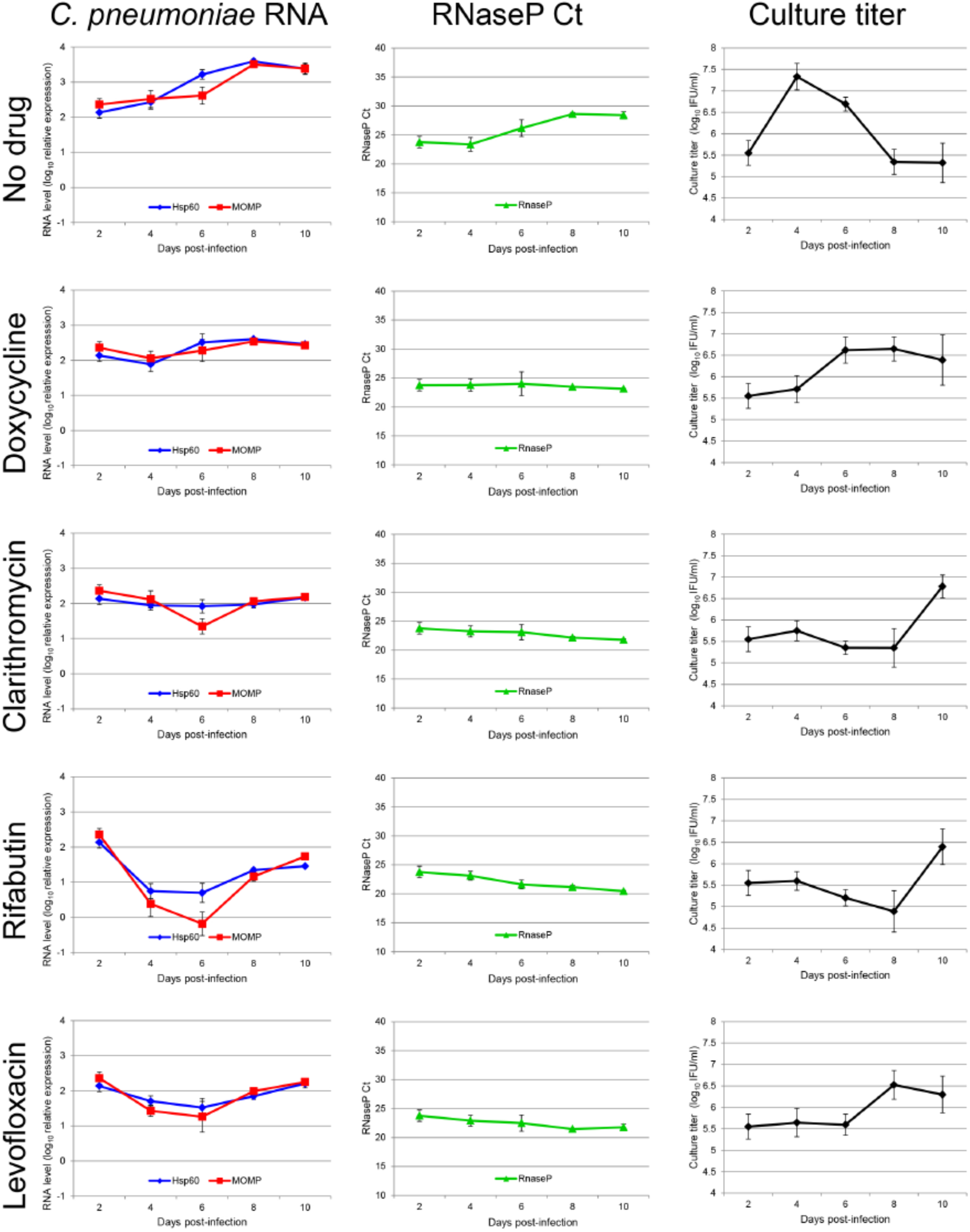
Evaluation of *C. pneumoniae* infectivity following antimicrobial treatment of established infection in HEp2 cells. HEp2 cells were infected at an MOI of 0.2 IFU/cell. Following two days of culture, one set of samples was collected for day 2 measurements and antibiotic free medium [No drug], 1 μg/ml doxycycline, 0.25 μg/ml clarithromycin, 0.25 μg/mlrifabutin, or 1 μg/ml levofloxacin was added to the remaining samples. At days 4, 6, 8, and 10 post-infection, samples wereeithercollectedorgivenantimicrobial-free medium. Expression of HSP60 and MOMP mRNA was measured in infected cell lysates using RT-qPCR. Results were normalized to RNAseP expression and are plotted relative to samples collected at day 0 post-infection. The RNAseP Ct values used to calculate RNA level are presented in the second column of graphs. Infectious titer was measured in infected cell lysates.

Relative expression of HSP60 and MOMP mRNA in untreated infection significantly increased at day 6 post-infection mainly due to declination in RNAseP expression, likely caused by unhealthy and dying cells undergoing unchecked *C. pneumoniae* replication. The decrease in infectivity following day 4 of infection supports this conclusion. In contrast, RNAseP levels remained constant or increased in cultures treated with antimicrobial agents, presumably protected from injurious effects of *C. pneumoniae* proliferation.

Post-treatment recovery of infections was also monitored by confocal microscopy to delineate morphologic features of inclusions that correlate with dynamics of RNA expression and infectivity [Fig 6]. Samples were infected and treated as outlined above. Inclusion diameters were measured over time, and a summary can be found in Table 1. At day 2 post-infection [time of treatment initiation], HSP60 staining demonstrated bright, consistently stained inclusions approximately 6.7 μm in diameter. By day 4 post-infection [two days of antimicrobial treatment], inclusions were brightly stained and significantly larger [14.2 ± 6.2 μm diameter], and more punctate compared to day 2 post-infection.

**Table 1.** Inclusion size following in vitro antimicrobial treatment and recovery in antimicrobial free Medium

|  | No drug | Doxycycline | Clarithromycin | Rifabutin | Levofloxacin |
| --- | --- | --- | --- | --- | --- |
| Day 2 | 6.8 ± 2.1 |  |  |  |  |
| Day 4 | 14.2 ± 6.2 | 8.0 ± 2.3 | 7.9 ± 2.7 | 7.3 ± 2.5 | 10.8 ± 3.9 |
| Day 6 | 5.0 ± 4.3 | 10.3 ± 4.9 | 8.4 ± 2.1 | 6.7 ± 2.0 | 12.7 ± 5.5 |
| Day 8 | 4.9 ± 4.3 | 8.9 ± 6.8 | 9.4 ± 3.3 | 7.7 ± 2.6 | 12.7 ± 6.1 |
| Day 10 | 7.3 ± 5.3 | 5.1 ± 3.7 | 14.8 ± 8.1 | 8.3 ± 4.3 | 5.2 ± 1.5 |

\*Diameter of inclusions measured in µm. n ≥ 21 for each experimental condition

**Fig 6.**
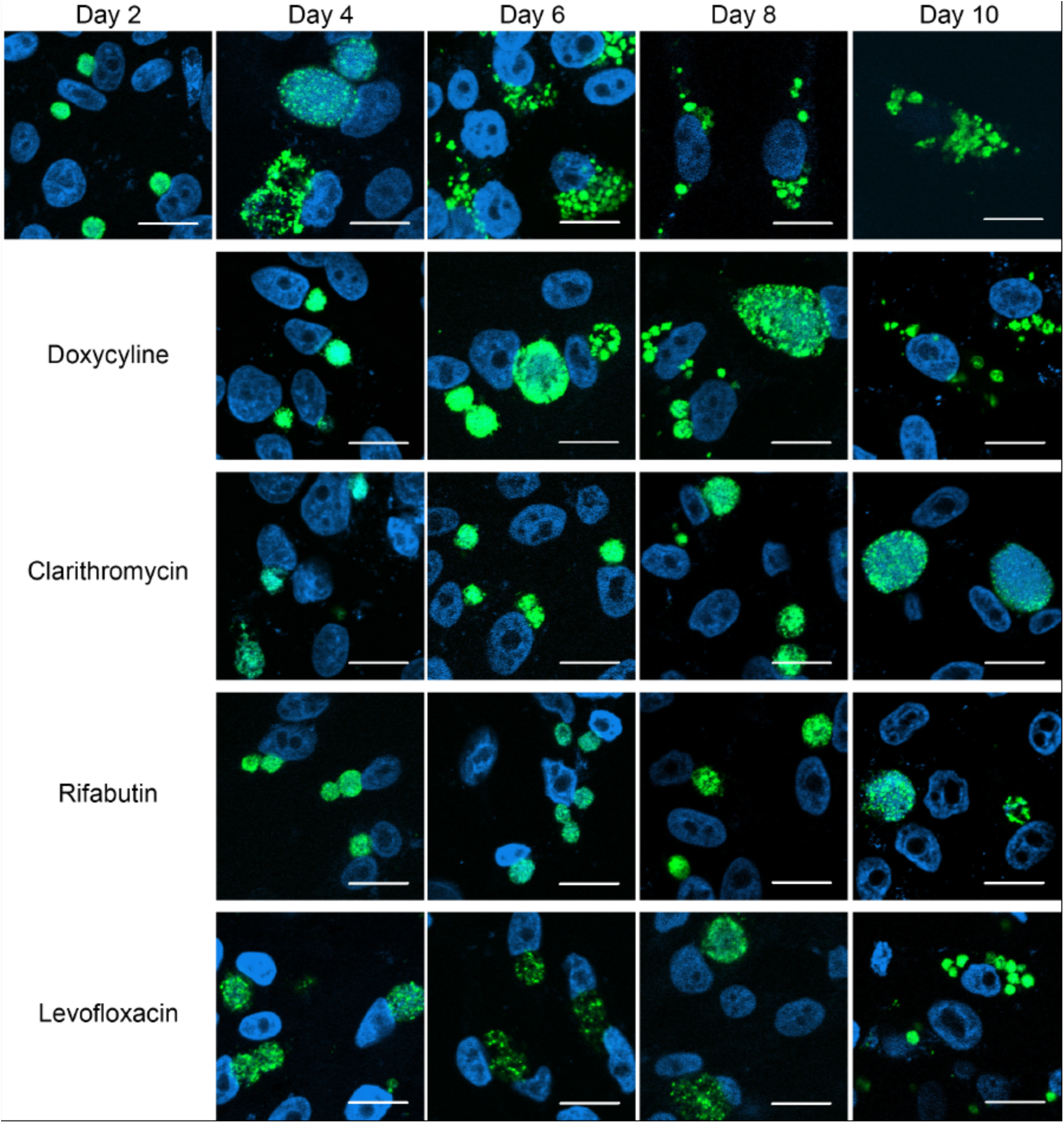
Inclusion morphology and infectivity of *C. pneumoniae* following antimicrobial treatment of established infection in HEp2 cells. HEp2 cells were infected at an MOI of 0.2 IFU/cell. Following two days of culture, one set of samples was collected for day 2 measurements and antimicrobial free medium[No drug], 1 μg/ml doxycycline, 0.25 μg/ml clarithromycin, 0.25 μg/ml rifabutin, or 1 μg/ml levofloxacin was added to the remaining samples. At days 4, 6, 8, and 10 post-infections, samples were either fixed or given antimicrobial-free medium. Fixed cells were stained with an anti-cHSP60 antibody [green] and ToPro3 [blue, DNA], and imaged using confocal microscopy. Scale bars, 20 μm.

Following day 4 post-infection, *C. pneumoniae*-mediated destruction of the cellular monolayer was observed in untreated cultures. HSP60 staining appeared as clusters of small inclusions or inclusion debris. Nuclear staining was diminished at day 8 and no longer visible at day 10. In doxycycline-, clarithromycin-, and rifabutin-treated infections, inclusions visualized at day 4 appeared very similar to those at day 2, with minimal increase in diameter. Inclusions in levofloxacin-treated infection were intermediate in size relative to day 2 and day 4 of the antimicrobial-free control culture. Removal of doxycycline corresponded to a pronounced increase in inclusion size at day 6 post-infection. Similar to untreated infection, *C. pneumoniae* appears to have overwhelmed the culture by day 8 post-infection, as clusters of inclusions and inclusion debris were present. In clarithromycin-treated infection, inclusion size moderately increased through day 8, followed by a more dramatic enlargement between day 8 and 10, from 9.6 ± 3.2 μm to 15.5 ± 7.4 μm. Inclusion size and morphology in the day 10 clarithromycin-treated culture were virtually indistinguishable from the day 4 untreated infection. In rifabutin-treated infection, significant increase in inclusion size was not observed until day 10 post-infection. At this time point, some inclusions resembled the day 4 pattern of untreated infection, though inclusions in the rifabutin-treated culture were smaller. In levofloxacin-treated infections,inclusions progressively increased in size and organization after removal of the antimicrobial agent, eventually evolving into a pattern characteristic of culture overtaken by infection. In general, an increase in infectivity was accompanied by an increase in inclusion size with exception of levofloxacin-treated infections in which inclusions were large until day 10 post-infection when the culture was overtaken by *C. pneumoniae*. The appearance of brightly stained, large inclusions correlated with an increase in infectivity in all antimicrobial treatment cell groups. With exception of rifabutin-treated infection, changes in mRNA levels were modest upon treatment and recovery. However, using the data provided by all three methods, we were able to detect recovery of *C. pneumoniae* replication after treatment with each antimicrobial agent. Furthermore, the kinetics and magnitude of recovery were dependent on the specific antimicrobial agent used.

## Discussion

In this pilot study, we determined MICs of doxycycline, clarithromycin, rifabutin, and levofloxacin for *C. pneumoniae* replication using a conventional susceptibility testing method based on detection of inclusion bodies as well as an RT-qPCR method for measurement of mRNA. The MICs were consistent, except for levofloxacin, with published results follows: doxycycline = 0.25 μ g/ml,clarithromycin = 0.0625 μ g/ml, rifabutin = 0.016 μ g/ml, and levofloxacin = 5μ g/ml. For levofloxacin, 1 μ g/ml was noted to be almost completely inhibitory, which was more in line with published results of 0.25 - 0.5 μ g/ml. Bactericidal activity for all four antimicrobial agents was achieved using a conventional susceptibility testing method based on chlamydial regrowth as well as by the RT-qPCR-based method. Evaluation of infectivity revealed no viable/infectious chlamydial elementary bodies in the day 3 cell lysates, and immunofluorescent confocal microscopy revealed no inclusions. However, if HEp2 cells after initial infection by *C. pneumoniae* elementary bodies had the antimicrobial agent removed and were then allowed to resume growth in fresh media over an additional three-day period, infectivity was noted for doxycycline, clarithromycin, and levofloxacin, but not for rifabutin. These results argue that only rifabutin is rapidly bactericidal in an initial infection of HEp2 cells by elementary bodies.

Moreover, the results after established infection was achieved in HEp2 cells was markedly different that results achieved in HEp2 cells after initial infection with elementary bodies. None of the antibiotics tested was inhibitory when these agents were added at day 2 post infection, although a reduction in the levels of mRNA was noted. The largest reduction of mRNA levels was seen with rifabutin [P<0.0001]. On day 4 post infection, an antibiotic-free medium exchange wasdone and the cultures were allowed to continue for a total of 10 days post-infection with fresh media added every two days. Infectivity under these conditions was noted to follow restoration of mRNA expression by approximately two days. In general, an increase in infectivity was accompanied by an increase in inclusion size by immunofluorescent confocal microscopy. The kinetics and magnitude of recovery were dependent on the specific antibiotic used, with clarithromycin and rifabutin having the slowest recovery at day 10 post-infection.

Other investigators (16-20) have noted that for *Chlamydia* spp., antimicrobial agents have a diminished effect if the antimicrobial agent is added after eight hours of incubation. This decreased activity of antimicrobial agents is consistent with the observations of these agents in a continuous-infection model (26-28), in which HEp2 cells were chronically infected with *C. pneumoniae* for longer than one year; 30 days of *in vitro* treatment of these cells with azithromycin, clarithromycin, and levofloxacin at concentrations comparable to those achieved in pulmonary tissue reduced, but did not eliminate *C. pneumoniae*. Results of this pilot study as well as these observations by other investigators suggest that further evaluation of susceptibility testing methods using a 48-hour established infection in cell cultures and RT-qPCR is needed to optimize treatment recommendations for chronic *C. pneumoniae* infections.

## Materials and Methods

### Cells, bacteria, and reagents

HEp2 cells [Quidel] were grown in DMEM containing 4.5 g/L glucose, 4.5 g/L L-glutamine, and 4.5 g/L sodium pyruvate [Mediatech] supplemented to contain 10% fetal calf serum [Seradigm], 100 μg/ml streptomycin [Sigma], and 0.25 μg/ml amphotericin B [Sigma]. *Chlamydia pneumoniae* strain AR-39 was purchased from ATCC and propagated as previously described (29). Antimicrobial agents considered to be active against *Chlamydia spp*. were tested; these included doxycycline, clarithromycin, levofloxacin, and rifabutin. All antimicrobial agents were purchased from Sigma.

### Quantification of chlamydial mRNA expression using RT-qPCR

Monolayers of HEp2 cells [∼4 × 10^5^ cells] seeded in 24-well plates were inoculated at an MOI of0.2 IFU/cell, and plates were centrifuged at 900 × g for 1h. After centrifugation, inoculum was removed, monolayers were washed with PBS [Mediatech], fresh medium supplemented with 1 μg/ml cycloheximide [Sigma] was added, and plates were incubated for various intervals at 35°C. Culture lysates were prepared by scraping cell monolayers into growth medium and were stored at -80°C. mRNA was isolated from cell lysates using the MagNA Pure LC RNA Isolation – High Performance Kit [Roche]. The mRNA HP Cells protocol was used with thefollowing modifications: 100 μl of lysate was pre-mixed with 100 μl lysis/binding buffer and mRNA was eluted in 100 μl elution buffer. Purified mRNA was stored at 4°C prior to RT-qPCR. The RT-qPCR reaction consisted of 10μl qScript SLT One step RT-qPCR Tough Mix [Quanta], 5 μl sample mRNA, 10 μM forward and reverse primers, and 10 μM Taqman fluorogenic probe in a total reaction volume of 20 μl. Target amplification, data acquisition, and data analysis were performed on an ABI Prism 7500 real-time PCR system [Applied Biosystems]. The cycling parameters were successively 15 min at 50°C, 5 min at 95°C, and 40 cycles consisting of 15 sec at 95°C and 32 sec at 60°C.ThefluorogenicprobesusedwereHSP60[GroEL1],5’-dFAM-ACACGCAACCATTGCAGCTGTCGA-BHQ-1-3	MOMP,5’-dFAM-ACTACTGCCGTAGATAGACCTAACCCGGCC-BHQ-1-3’;andRNAseP,5’-dFAM-TTCTGACCTGAAGGCTCTGCGCG-BHQ-1-3’[BiosearchTechnologies].TheprimersusedwereHSP60F,5’TGAAAGAGAAAAAAGACAGAGTAGATGATG3’;HSP60R,5’GTTCCACCACCAGGGAGGAT3’;MOMPF,5’GGATCCGCTGCTGCAAACTA-3’;MOMP-R,5’-CACTCTGCATCGTGTAAATGCTT-3’;RNAseP-F,5’-AGATTTGGACCTGCGAGCG-3’;andRNAsePR,5’GAGCGGCTGTCTCCACAAGT-3’[Life Technologies]. Amplification efficiency for each primer/probe set was experimentally determined to be 92.3%, 94.4%, and 93.7% for HSP60, MOMP,and RNAseP, respectively. mRNA expression was calculated using the ΔΔCt method (30).

### Assay for *C. pneumoniae* infectivity

Lysates of *C. pneumoniae*-infected HEp2 cells were prepared as described above, sonicated on ice [10-15 watts for ∼ 5 sec] and serially diluted in PBS. Lysate dilutions were used to inoculate fresh HEp2 cells growing on glass coverslips [Electron Microscopy Sciences] treated with poly-L-lysine [R&D Systems]. Cultures were fixed with methanol at 72 h post-infection and stained with a fluorescein-conjugated murine monoclonal antibody to *Chlamydia* [genus-specific] [Biorad]. Coverslips were mounted on slides using Aqua Poly/Mount [Polysciences, Inc], and IFU/ml was determined using an Olympus BX41 fluorescence microscope. The limit of detection of this assay is 134 IFU/ml [2.1 log_10_ IFU/ml].

### Confocal immunofluorescence microscopy for detection of *C. pneumoniae* inclusions

HEp2 cells growing on glass coverslips in 24-well plates were infected as described above, fixed at various intervals, and stained with a mouse cHSP60-specific monoclonal antibody [Acris Antibodies], followed by Alexa 488-conjugated anti-mouse secondary antibody and ToPro3 [Thermo Fisher Scientific]. Images were acquired using a Zeiss LSM 510 META inverted confocal microscope. Inclusion diameters were measured using LSM Image Browser software [Zeiss].

### Statistics

Results are expressed as means ± SD. One-way ANOVA and Tukey’s multiple comparison tests where calculated using Prism 4 for Windows [Graphpad]. *P* values less than 0.05 were considered statistically significant.

## Acknowledgements

Acquisition and analysis of confocal microscopy images were made possible in part through use of the Vanderbilt University Medical Center Imaging Shared Resources. Financial support was provided by gifts from Paul Griffin, Ursula Hess, and David Wheldon.

